# Mitochondrial double-stranded RNA accumulation in brain aging and Alzheimer’s disease

**DOI:** 10.64898/2026.02.02.703345

**Authors:** Rachel L. Doser, Thomas J. LaRocca

## Abstract

Mitochondria and inflammation are tightly linked in aging and Alzheimer’s disease (AD), and recent evidence implicates mitochondrial double-stranded RNA (mt-dsRNA) as a potential trigger of inflammation. We examined mt-dsRNA accumulation and dsRNA signaling in brain aging and AD using human brain tissue and complementary *in vitro* transcriptomic datasets, quantifying mitochondrial transcripts and dsRNA editing. We found that mt-dsRNA accumulated after midlife and coincided with reduced expression of mitochondrial RNA processing and translation machinery, along with increased expression of dsRNA antiviral signaling proteins, consistent with cytoplasmic mt-dsRNA-driven inflammation. In AD brains, mt-dsRNA accumulation was further increased and correlated with cognitive impairment, neuropathological severity, and AD risk genotypes. Genes associated with these measures reflected altered ubiquitin-dependent regulation of antiviral signaling, potentially indicating altered sensitivity to mt-dsRNA. Together, these findings highlight mitochondrial RNA homeostasis as an unrecognized contributor to age- and AD-related neurodegeneration by identifying mt-dsRNA as a potential driver of chronic inflammation.

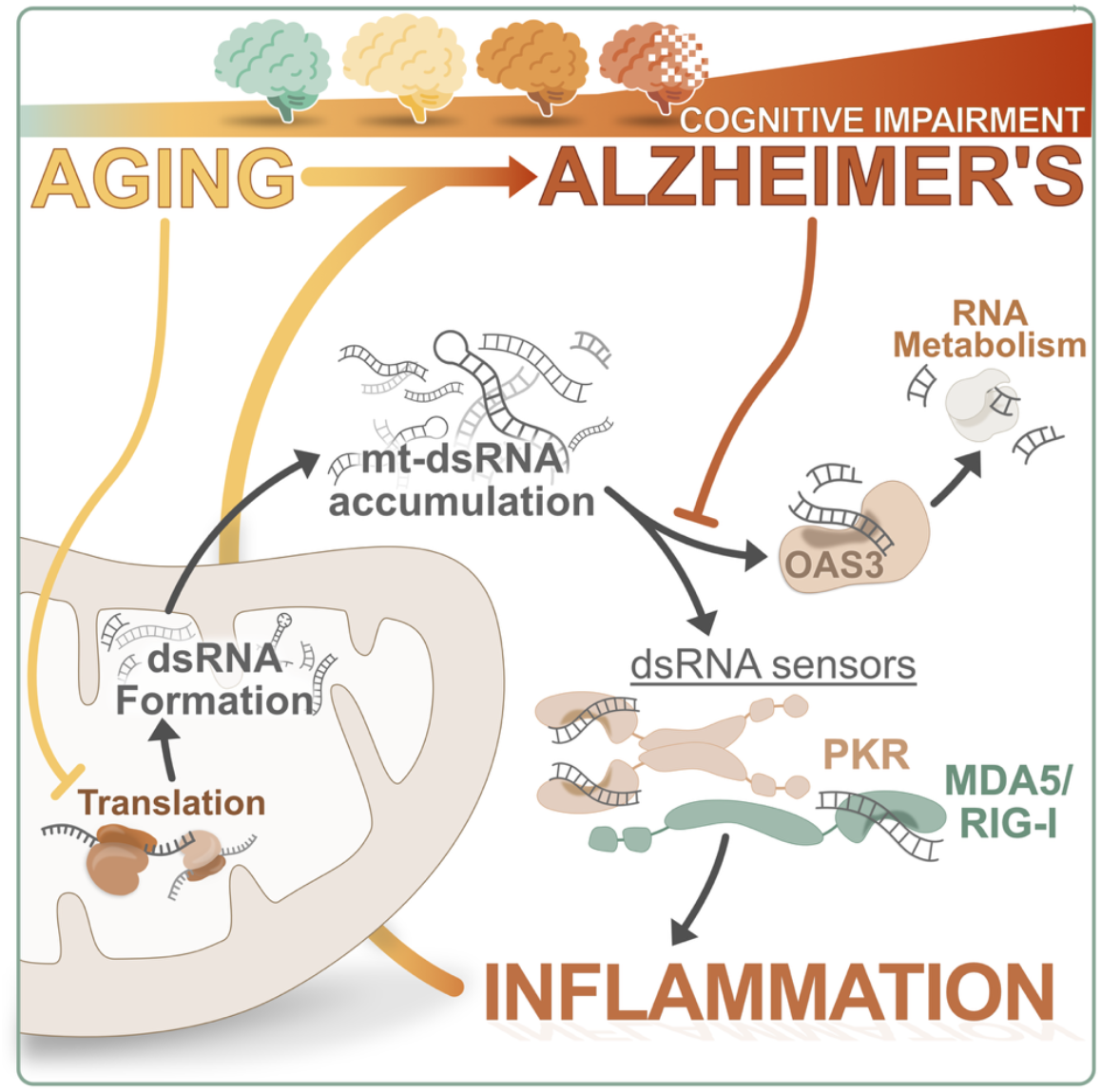

## INTRODUCTION

Aging is the leading risk factor for Alzheimer’s disease (AD), and understanding age-related molecular processes that increase AD susceptibility remains a central challenge in the field^1^. Among the cellular hallmarks of aging and AD, mitochondrial dysfunction is especially prominent and encompasses impairments in mitochondrial metabolism, proteostasis, and quality control mechanisms such as fission, fusion, mitophagy, and biogenesis^2–4^. Historically, studies have focused on impaired oxidative phosphorylation and mitochondrial dynamics as consequences of these impairments and drivers of neuronal dysfunction and cell death in AD. However, recent evidence suggests that dysregulation of the mitochondrial transcriptome may also be a consequence of mitochondrial dysfunction that contributes to important age- and disease-related processes^5^, including inflammation.

Transcription of the mitochondrial genome occurs bidirectionally, producing long polycistronic transcripts that require extensive post-transcriptional processing, including cleavage, polyadenylation, RNA editing, and folding, to generate functional messenger RNA^6,7^. Proper coordination between mitochondrial translation and RNA processing is essential to mitochondrial gene expression, and to prevent accumulation of homologous transcripts^8,9^. When this coordination fails, unprocessed or overly abundant RNAs can form double-stranded RNA (dsRNA) within mitochondria (mt-dsRNA)^9,10^. Recent studies have shown that mt-dsRNA can be released into the cytoplasm under both stress and physiological conditions^11^. Because dsRNA is a molecular pattern typically associated with viral infection, it is a potent immune activator; as such, once in the cytoplasm, mt-dsRNA can promote sterile inflammation by binding to pattern recognition receptors such as RIG-I/DDX58, MDA5/IFIH1 and PKR/EIF2AK2, thereby triggering type I interferon (IFN) signaling^12,13^. Because mitochondrial dysfunction occurs with aging and AD, these recent findings suggest the possibility that mt-dsRNA may be an unrecognized driver of sterile, chronic inflammation, which is a central feature of brain aging and AD^12,14–16^.

Despite the above observations, mt-dsRNA has not been systematically examined in the context of brain aging or AD. Therefore, in the present study we set out to: 1) profile mitochondrial RNA dynamics with age, 2) determine if mt-dsRNA accumulation occurs in aging and/or AD, and 3) test the idea that these events might be related to cognitive decline and AD risk. To accomplish this, we analyzed transcriptomic data from two large postmortem brain datasets to characterize changes in mitochondrial RNA expression and dsRNA footprints in the aging brain, and in individuals with AD. Our analyses suggest that the mitochondrial transcriptome is relatively unchanged with age, but the expression of important mitochondrial RNA processing machinery declines with age—and these changes are accompanied by elevated mt-dsRNA signatures that correlate with innate immune activation. Mitochondrial transcripts and evidence of mt-dsRNA accumulation are also increased in AD brains and related to cognitive decline and genetic risk for AD. Together, our findings suggest that failure to properly regulate mitochondrial RNA homeostasis may lead to the accumulation of immunogenic mt-dsRNA contributing to neuroinflammatory signaling in aging that is worsened in AD.

## METHODS

### RESOURCE AVAILABILITY

#### Lead Contact

Requests for further information and resources should be directed to the Lead Contact and Senior Author, Thomas LaRocca.

#### Materials Availability

This study did not generate new unique reagents.

#### Data and Code Availability

All RNA-seq datasets used in this study are publicly available. The main datasets used here for analyses are total RNA-seq data from human prefrontal cortex and include: the NABEC dataset (105 neurologically normal individuals aged 19–86 years; see Table S1; Synapse ID: syn3270007) and ROSMAP dataset (218 individuals aged >73 years with clinical diagnoses of no cognitive impairment [NCI], mild cognitive impairment [MCI], or AD [sporadic or genetic]; see Table S2; Synapse ID: syn3219045). Supporting RNA-seq datasets include: PKR immunoprecipitated RNA in HeLa cells (GSE108986)^17^, directly reprogrammed/induced human neurons (E-MTAB-3037)^18^, DRP1 in human neuronal co-culture (GSE237013)^19^, human muscle biopsy following *in vivo* Urolithin A treatment (GSE197273)^20^, and BAX/BAK knock-out in human lung fibroblasts (GSE196610)^21^. Processed data tables and normalized mt-dsRNA editing values are available from the corresponding author upon request, and analysis scripts can be found on GitHub at: racheldoser/Mitochondrial-dsRNA-RNAseq-analyses^22^.

## METHOD DETAILS

### RNA-seq Data Processing and Normalization

Raw sequencing reads were aligned to the human reference genome (GRCh38) using STAR (v2.7.3a) with default settings (Dobin et al., 2013). Mitochondrial transcript abundance was quantified as total reads mapping to ChrM, normalized by library size to generate sample-specific scaling factors.

### mt-dsRNA Signature Quantification

A-to-I RNA editing events, a footprint of cytoplasmic dsRNA^23^, were identified using the SPRINT (SNV calling pipeline for RNA editing identification) workflow^24^, which provides high-confidence editing calls by filtering out single nucleotide variations, sequencing errors, and mapping artifacts. Because the total RNA-seq libraries in the two main datasets analyzed were generated using rRNA depletion approaches, mt-rRNAs observed in corresponding analyses likely represented post-depletion residual transcripts, which are common and substantial in most datasets. To account for potentially differences in depletion efficiency, samples with drastically low or high ChrM sequencing depth were excluded from our analyses which was done by including only samples whose ChrM scaling factor fell within the 5^th^ to 95^th^ percentile. The total number of editing counts from all ChrM transcripts was also normalized to each sample’s ChrM scaling factor (to further account for sample-specific variation in mitochondrial rRNA depletion efficiency by scaling editing counts to overall mitochondrial transcript abundance), thereby providing a more conservative estimate of mt-dsRNA-associated editing across samples. These normalized A-to-I editing values were used in all downstream correlation and group comparison analyses. Population distribution of ChrM transcript edits by location/gene was performed by binning raw edits every 200 nucleotides for all age groups in the NABEC dataset.

### Differential and Relative Gene Expression Analyses

Differential gene expression analyses were conducted on raw gene counts using DESeq2 (v1.30.1)^25^. Where indicated, additional variables (e.g., APOE genotype) were controlled for by accounting for the variable in the design function (i.e., design = ∼APOEgenotype + condition). Genes with an adjusted p-value (FDR) <0.05 were considered significantly differentially expressed. To calculate relative expression of a subset of genes, the average normalized expression of a gene for a given group was divided by either average normalized expression in younger (<35 years, for NABEC) or NCI subjects (for ROSMAP). Results were FDR-corrected for multiple testing unless otherwise noted.

### Gene Co-expression and Network Analysis

Weighted gene co-expression network analysis (WGCNA) was performed on the ROSMAP dataset using the WGCNA R package (v1.70-3)^26^. Expression data were filtered to include only protein-coding genes. A signed co-expression network was constructed using a soft-thresholding power of 9, and modules were identified using dynamic tree cutting with a minimum module size of 300 genes. Closely related modules were merged using a merge cut height of 0.35. Module eigengenes were calculated and correlated with clinical traits, including MMSE scores, APOE genotype, Braak (tau) and CERAD (amyloid plaques) pathology, and mt-dsRNA signatures. Enrichment for mitochondria-relevant genes was evaluated using the MitoCarta 3.0 database^27^ and the GeneOverlap R package^28^.

### Correlation Analyses

Correlation analyses were conducted on normalized data. Only genes with reads in more than half of the subjects were retained. Pearson product-moment correlation coefficients were calculated using JMP Pro (version 18) between mt-dsRNA signatures or clinical measures (e.g., MMSE or Braak Score) and gene lists (e.g., all protein-coding or those involved in dsRNA signaling or mitochondrial RNA homeostasis). Correlation matrices were visualized with heatmaps, and significance was assessed using the correlation probability.

### Enrichment Analyses

For functional enrichment analyses, gene sets overlapping between cognitive decline (MMSE score), tau pathology (Braak stage), and mt-dsRNA levels were subjected to Gene Ontology (GO) analysis using the NIH DAVID Bioinformatics Resources (v6.8)^29^. All GO terms with a raw p-value <0.05 were exported and submitted to ReVIGO^30^ to remove redundant terms and cluster similar biological processes using semantic similarity. Fold enrichment values and representative GO terms were used for visualization. Enrichment for mitochondria-relevant functions was assessed via a hypergeometric overlap (using GeneOverlap R package)^28^ between gene lists and MitoCarta 3.0 pathways^27^.

### Statistical Analysis

Statistical analyses were performed using either Prism5 software (GraphPad Software), R Studio, or JMP Pro Version 18 (see methods sections above for more detail). Generally, group comparisons were evaluated using ordinary one-way ANOVA with correction for multiple comparisons (Sidak) where applicable. Correlation significance was determined by Pearson’s r test and enrichment p-values were computed via hypergeometric testing. Adjusted p-values (FDR or Bonferroni adjusted [P_adj_]) were considered statistically significant using an α < 0.05. Bar plots represent mean ± SEM. The distribution of each dataset was assessed for normality using a Shapiro-Wilk test. All mt-dsRNA signature datasets were non-normally Gaussian distributed, so Brown-Forsythe and Welch ANOVA tests were used to compare mt-dsRNA editing between groups with a Dunnet T3 correction for multiple comparisons.

### Declaration of AI-assisted technologies in the writing process

During the preparation of this manuscript, the authors used ChatGPT only to proofread text and to enhance readability and language of the manuscript. Generative AI was not used to create or modify any data or figures in any way. After using this tool/service, the text was re-reviewed and edited as needed to ensure the content was correct and representative of the presented findings. Authors acknowledge that they are responsible for the article’s content despite AI assistance with text editing.

## Acknowledgements

The authors would like to thank Drs. Chris Link (University of Colorado Boulder) and Karyn Hamilton (Colorado State University) for their input on this topic and manuscript.

## Author Contributions

R.D and T.L. performed all bioinformatics analyses. R.D. constructed the figures. R.D. and T.L. wrote and edited the manuscript.

## RESULTS

### Accumulation of mitochondrial dsRNA in brain aging

To understand whether the mitochondrial genome (ChrM) may be an unrecognized source of dsRNA during brain aging, we analyzed RNA-seq data from post-mortem human brain tissue of neurologically normal individuals (NABEC dataset; n = 105, age 19–86 years; Table S1). First, we quantified the expression of the mitochondrial genome (sum of reads for all ChrM-encoded transcripts) across the lifespan and found that ChrM expression remained relatively stable in the brain throughout aging (Figure 1A). We also compared the expression of all genes, both nuclear- and ChrM-encoded, in older (>56 years) versus younger (<35 years) individuals (Figure 1B) to identify differentially expressed genes (DEGs) in older brains (Age-DEGs). We found that most ChrM transcripts were expressed at similar levels. However, many transcripts for nuclear-encoded genes that localize to and aid in the function of mitochondria (MitoCarta genes)^27^ were either increased (32 genes) or decreased (40 genes) in older brains (Figure 1B, green points), whereas transcripts encoding mitochondrial rRNAs were decreased in the older individuals (MT-RNR1: FDR=0.025; MT-RNR2: FDR=0.055; Figure 1B, red points). A gene set enrichment analysis on differentially expressed MitoCarta transcripts indicated that genes related to folate, one-carbon, reactive oxygen species (ROS) and glutathione metabolism, and mitochondrial dynamics and surveillance were enriched in the increased Age-DEGs. In contrast, genes encoding key players in mitochondrial respiration, lipid metabolism and translation were enriched in the reduced Age-DEGs (Figure 1C). In line with this finding, many genes important for local translation of mitochondrial RNAs (mtRNAs), specifically those important for ribosome function, were generally downregulated in older age groups compared to individuals <35 years of age (Figure S1A and S1B). Expression of genes encoding mtRNA processors was also slightly increased, and those related to metabolism were further elevated with older age (Figure S1A and S1B).

**Figure 1.**
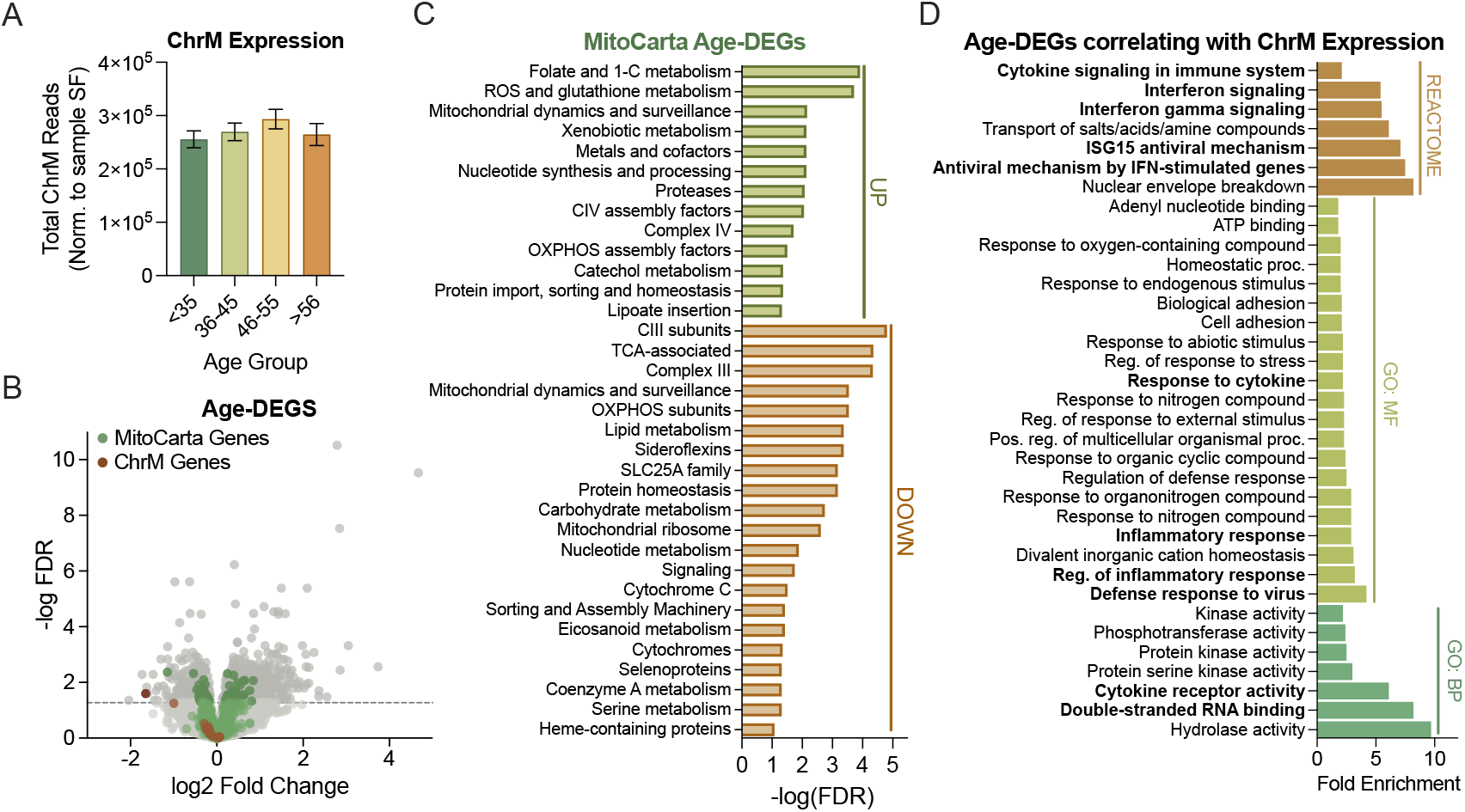
Reduced mitochondrial RNA processing and translation in the brain with aging may lead to accumulation of double-stranded RNA (mt-dsRNA). A-F) Analyses of total RNA in the brain from individuals 19-86 years old (NABEC dataset; n=105). A) Total RNA transcripts from the mitochondrial chromosome (ChrM) normalized to library size for each sample (size factor [SF]). B) Volcano plot of differentially expressed genes in older (>55 years) compared to younger (<35 years) individuals (Age-DEGs). Green points indicate nuclear-encoded MitoCarta genes and red points indicate ChrM transcripts. Dark grey/green/red points above the dashed line represent genes that are significantly differentially expressed at older ages (FDR < 0.05). C) Gene set enrichment analysis of up- or down-regulated AgeDEGs and MitoCarta 3.0 pathways. D) KEGG REACTOME, gene ontology biological processes (GO:BP) and molecular function (GO:MF) for Age-DEGS significantly correlated (adj. p-value <0.05) with ChrM expression.

Although ChrM expression remained consistent with aging, the above results suggested that local mitochondrial translation may decline in the aged brain, potentially causing mtRNA (and perhaps mt-dsRNA) accumulation. Consistent with this idea, we noted that the Age-DEGs whose transcript levels correlated with ChrM expression were largely related to dsRNA sensing and type-I immune activation (Figure 1D). Similar to others’ findings^31–33^, the expression of genes related to dsRNA and type-I interferon (IFN) signaling was generally greater in older brains (Figure S1C). Because existing data indicate that dsRNA sensing occurs in the nucleus and cytoplasm^34^, mt-dsRNA would need to be released from mitochondria to initiate such dsRNA/IFN signaling. Again, this possibility, the expression of some non-selective mitochondrial pores/channels (e.g., BAX/BAK channels) was increased in older brains (Figure S1D). However, to more directly estimate the levels of this putative mt-dsRNA, we turned to the SNP-free RNA editing Identification Toolkit (SPRINT)^35^, a bioinformatics pipeline that quantifies adenosine-to-inosine (A-to-I) editing, which is a dsRNA “footprint” that occurs only in cytoplasmic or nuclear dsRNA^23^ due to the predominant localization of dsRNA editing enzymes (namely adenosine deaminase acting on RNA [ADAR] enzymes) to these subcellular compartments^36^. Our quantification of mtRNA A-to-I editing revealed that nearly all ChrM transcripts were edited, and some transcripts (e.g., RNR1 and RNR2) had proportionally more A-to-I edits (Figure 2A).

**Figure 2.**
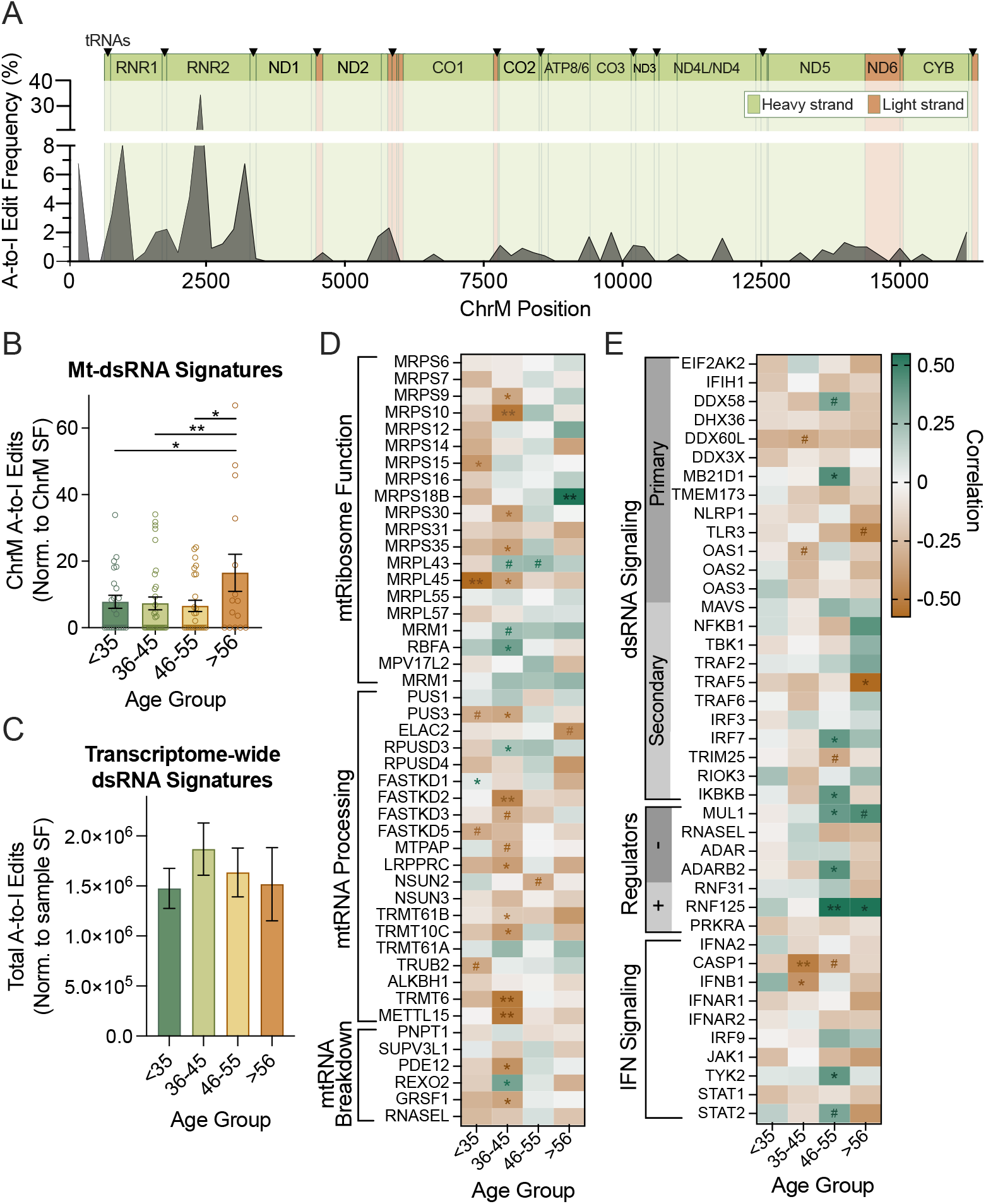
Decreased RNA processing and translation at older ages may cause mt-dsRNA formation and upregulation of mitochondria-localized dsRNA immune sensors. A-E) Analyses of total RNA from brains of individuals 19-86 years old (NABEC dataset; n=105). A) Frequency of A-to-I edits across ChrM (binned every 200 bp; green regions: H-strand encoded, orange regions: L-strand encoded). B) Quantification of A-to-I edits in mt-RNA transcripts normalized to ChrM size factor (ratio of total ChrM reads for a sample divided by dataset mean) for each age group. C) Quantification of transcriptome-wide A-to-I edits (normalized to sample SF) in each age group. D and E) Correlations among mt-dsRNA signatures (A-to-I edits) and expression of genes relevant for (D) mitochondrial RNA processing, metabolism and translation or (E) dsRNA immune signaling. *:p<0.05, **:p<0.005, ***:p<0.001.

Although the SPRINT pipeline was designed to distinguish edits from single nucleotide variations (such as those that may arise due to ChrM mutations), we analyzed the presence of homologous reads, secondary structure and editing sites of the most abundantly edited ChrM transcript, MT-RNR2, to verify the pipeline’s mtRNA editing detection. We found that MT-RNR2 may readily form intra-strand dsRNA (Figure S2A) and on average, 17.7% of detected MT-RNR2 reads are antisense and thus homologous to coding transcripts (suggesting the possibility of inter-strand dsRNA). Additionally, the majority of A-to-I editing sites (67.8%) were edited in more than one subject, demonstrating the pipeline’s ability to distinguish edits from nucleotide variants. These pieces of information suggest that mtRNA can form dsRNA from both intra- and inter-strand interactions, and that mt-dsRNA can be detected via quantification of A-to-I edits with the SPRINT pipeline. However, to further confirm this idea, we analyzed RNA immunoprecipitation sequencing data from experiments targeting native dsRNA binding proteins (e.g., Protein Kinase R [PKR]). These analyses also indicated that mt-dsRNA is formed and released under non-pathological contexts. In fact, mtRNAs were among some of the most enriched transcripts in PKR co-IP compared to total RNA pools (Figure S2B). Furthermore, to assess whether A-to-I edited mt-RNA was representative of cytoplasmic mt-dsRNA bound to PKR, we tested the relationship between transcript A-to-I editing and enrichment in the PKR co-IP. The majority (89.2%, p=2.76×10^-6^) of A-to-I edited mt-dsRNA was immunoprecipitated with PKR compared to total RNA input, further supporting our methodology for mt-dsRNA estimation.

After this technical validation, we next quantified mt-dsRNA signatures (A-to-I edits) in the NABEC dataset to more directly assess mt-dsRNA transcripts in the brain across the lifespan. We found that mt-dsRNA signatures remained stable until ∼55 years of age, after which the abundance of A-to-I edited ChrM transcripts increased two-fold (Figure 2B). We analyzed A-to-I editing for the entire transcriptomes and found that dsRNA editing is relatively consistent in the brain with aging (Figure 2C) indicating that mtRNA may indeed be particularly predisposed to formation of dsRNAs. Mt-dsRNA accumulation appeared to be innate to aging cells, as this finding was recapitulated in a separate *ex vivo* dataset on human fibroblast-derived induced neurons from subjects between the ages of 30-89 (Figure S2C). Importantly, in the original report on these induced neurons, cells from aged donors maintained aging phenotypes including mitochondrial dysfunction^37^. Therefore, to gain insight into whether mitochondrial dysfunction may contribute to mt-dsRNA formation and release at older ages, we analyzed other publicly available RNA-seq experiments. In one dataset, we found that inhibition of mitochondrial fission (with P110) in co-cultures of human neurons and glial cells resulted in a slight, although non-significant increase in mt-dsRNA signatures (32%, p=0.5344; Figure S2D). In a second dataset, we found that upregulating mitophagy (degradation of damaged/dysfunctional mitochondria) in humans with the small molecule Urolithin A caused trending reduction in ChrM A-to-I edits in muscle tissue (46%, p=0.1727; Figure S2E). Finally, in another dataset we found that genetic knock-out of BAX/BAK in human lung fibroblasts prevented mt-dsRNA release (based on the absence of A-to-I edits) following X-ray-induced senescence and mitochondrial dysfunction in 2 out of 3 experimental replicates (Figure S2F). Taken together, these observations in human brains/neurons and proof-of-concept analyses of other tissues/datasets support the idea that age-related declines in mitochondrial homeostasis may contribute to the accumulation of mt-dsRNA.

### Potential mechanisms of mt-dsRNA accumulation and its consequences in brain aging

To understand what may occur in brain mitochondria that could lead to mt-dsRNA accumulation when homeostatic mechanisms decline with aging, we examined correlations among mt-dsRNA signatures and the expression of genes involved in mitochondrial RNA metabolism and translation (NABEC dataset; Figure 2D). In younger brains (<35 years), expression of many mitochondrial ribosomal proteins, including MRPS and MRPL family members was generally negatively related to mt-dsRNA signatures, but this was reversed in older age groups (Figure 2D). We also noted that: 1) expression of RNA processing genes, such as FASTKD and TRMT family members, was generally negatively related to mt-dsRNA signatures across the lifespan; and 2) gene expression related to RNA breakdown in mitochondria was generally anti-correlated with mt-dsRNA signatures, except in the 46-to-55-year-old age group (Figure 2D). Collectively, these associations suggest that, in addition to changes in mitochondrial function, age-related declines in mitochondrial RNA handling may contribute to mt-dsRNA accumulation.

Next, to investigate the relationship between age, cytoplasmic mt-dsRNA accumulation, and immune signaling, we examined correlations among mt-dsRNA signatures and the expression of dsRNA sensors, as well as related secondary immune signaling proteins and modulators across age groups (Figure 2E). Mt-dsRNA signatures were negatively related to most dsRNA sensors across the lifespan but positively correlated with DDX58/RIG-1 and MD21D1/cGAS in 46-to-55-year-old brains. In contrast, trending positive correlations were present for mt-dsRNA signatures and expression of most secondary dsRNA signaling proteins (MAVS, NF-κB, IRF7 and IKKβ) in the oldest age group. Expression of two dsRNA signaling modulators, MUL1 and RNF125, also positively correlated with mt-dsRNA signatures at older ages (>46 years), while other dsRNA modulators (e.g., RNASEL, RNF31 and ADARs) were negatively related to dsRNA signatures in older age. Some genes involved in IFN signaling, specifically those in the JAK/STAT pathway (which mediates dsRNA-triggered inflammation and related apoptosis), were also slightly or significantly positively correlated with mt-dsRNA signatures at older ages (Figure 2E). Interestingly, despite general increases in IFN gene expression with aging (Figure S1C), expression of many key IFN genes (e.g., CASP1 and IFNB) were inversely correlated to mt-dsRNA signatures in middle and older age groups (Figure 2E) suggesting that accumulation of mt-dsRNA may occur during instances of immunosuppression.

### Alzheimer’s disease-related changes in mt-RNA

Age-related mt-dsRNA accumulation and inflammation may be particularly important in AD pathogenesis for a few reasons: 1) abnormal mitochondrial maintenance and function is one of the earliest detectable changes in AD; and 2) antiviral defenses are impaired with mild cognitive impairment (MCI) and AD^38,39^, which may upregulate dsRNA signaling in these conditions^40^. To determine if mt-dsRNA accumulation in older brains is worsened with AD, we analyzed total RNA-seq data from dorsolateral prefrontal cortex samples of individuals with no cognitive impairment (NCI), MCI, or AD (ROSMAP cohort; >78 y.o.; n=218; Table S2). First, we found that ChrM transcript abundance was slightly (but not significantly) decreased in MCI compared NCI controls, and this change was nearly significantly reversed (p = 0.0759) in AD compared to MCI (Figure 3A). Next, we identified AD-related differentially expressed genes (AD-DEGs) and found that the two ChrM-encoded rRNAs, MT-RNR1 and MT-RNR2, were the most upregulated genes in AD brains (Figure 3B). Other mitochondrial genes, such as MT-ND3 and MT-CO2, were downregulated in AD relative to NCI, indicating that ChrM expression is not globally up- or downregulated with AD. Surprisingly, nuclear-encoded genes important for mitochondrial function (MitoCarta genes) were not differentially expressed in AD brains in this analysis (data not shown) suggesting that mechanisms by which ChrM expression and mt-RNA homeostasis are altered in AD may differ from those involved in aging.

**Figure 3.**
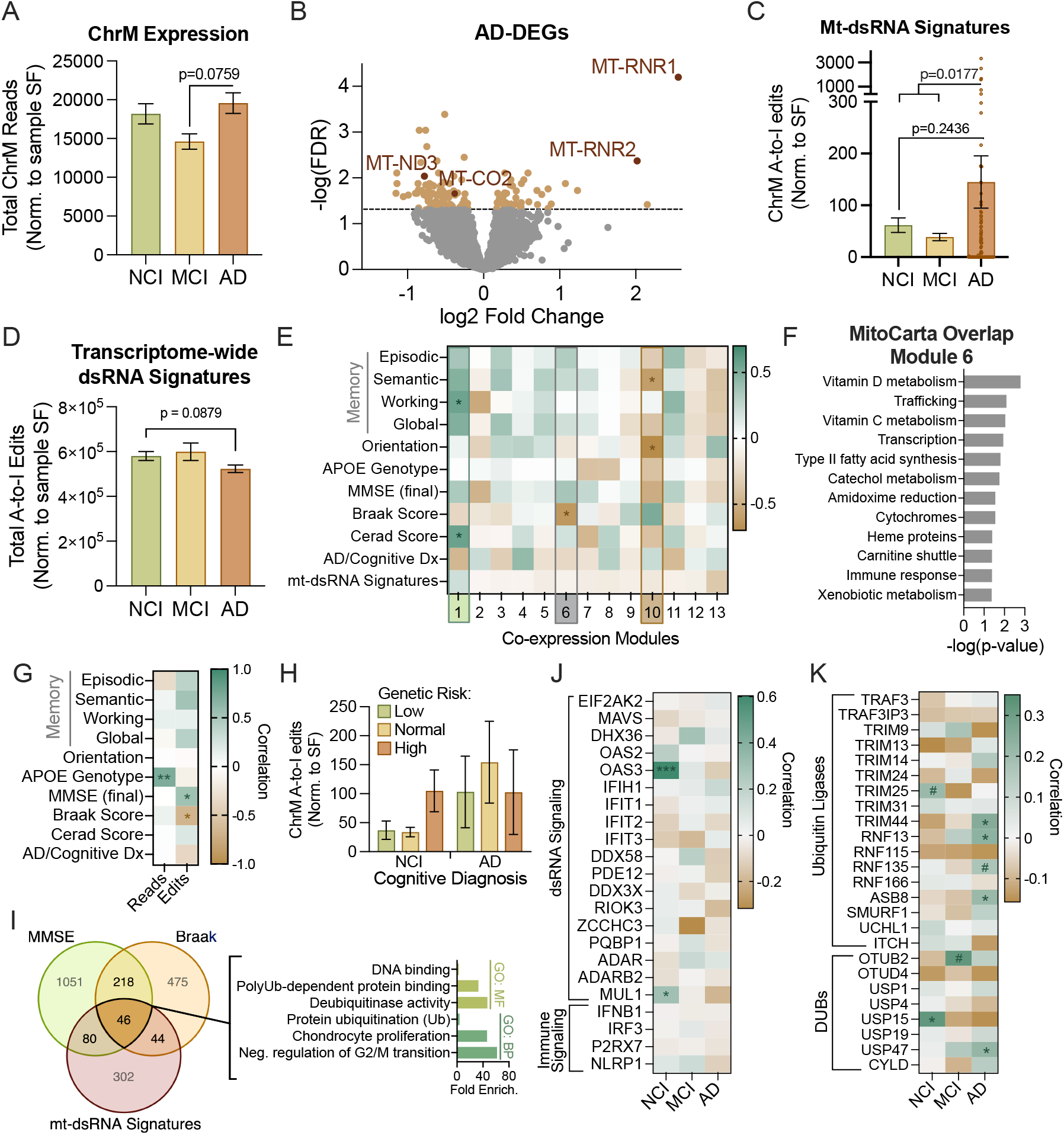
Increased mitochondrial genome expression and mt-dsRNA in Alzheimer’s disease correlates with pathology, genetic risk and cognitive dysfunction. A-K) Analyses of total RNA in the brain from individuals >73 years old with no cognitive impairment (NCI), mild cognitive impairment (MCI), or Alzheimer’s disease (AD; ROSMAP dataset; total n=218). A) Total RNA transcripts from the mitochondrial chromosome (ChrM) normalized to library size for each sample (size factor [SF]). B) Volcano plot of differentially expressed genes in AD compared to NCI subjects (AD-DEGs; orange data points FDR < 0.05; red data points = ChrM transcripts with FDR < 0.05). C) Quantification of A-to-I edits in mt-RNA transcripts normalized to ChrM SF (total ChrM reads for a sample divided by dataset mean) for each group. D) Quantification of transcriptome-wide A-to-I edits (normalized to sample SF) in each group. E) Weighted Gene Network Correlation Analysis (WGCNA) of protein-coding genes from a subset of individuals with extensive clinical measures of cognitive function (including memory tests and mini-mental state exams [MMSE]), AD risk genotype (copies of APOE4 allele) and quantification of AD pathology (Braak [severity of neurofibrillary tau tangles] and CERAD [neuritic plaque density]). Gene expression module 6 (grey box) contains the majority of ChrM transcripts. *:Padj<0.05. F) MitoCarta processes in which module 6 genes are enriched (using a hypergeometric overlap between gene lists). G) Pearson’s correlation of clinical traits with ChrM expression (reads) or mt-dsRNA signatures (edits) within a subset of ROSMAP samples (n=14) on which clinical information was available. H) Quantification of mt-dsRNA signatures in individuals with low (at least one copy of APOE2), normal (heterozygous for APOE3) or high (at least one copy of APOE4) genetic risk of AD based on APOE alleles. I) Left - Overlap of protein-coding genes that correlate with cognition (MMSE score), tau pathology (Braak score) and/or mt-dsRNA signatures. Right - Functional annotation of genes common to all three correlations. J and K) Pearson correlations among mt-dsRNA signatures and: J) genes involved in dsRNA signaling and related immune response or K) general and ubiquitin-dependent regulation of RIG-1/MAVS signaling. #: p<0.1, *:p<0.05, **:p<0.005, ***:p<0.001 (correlation probability).

To understand what might underlie AD-related differences in gene expression, we performed functional analyses of AD-DEGs and found that upregulated AD-DEGs were related to RNA modifications, DNA damage response, immune signaling involving nuclear factor kappa B (NF-κB) and mitogen-activated protein kinase (MAPK), and ubiquitin-dependent protein degradation (Figure S3A). Downregulated AD-DEGs were enriched for functional pathways related to epigenetic regulation, metabolism, and cell proliferation/differentiation (MAPK1/MAPK3 signaling; Figure S3A). These gene expression patterns are in line with what others have reported in AD^39,41^, and they could reflect both causes or results of mitochondrial dysfunction (including dsRNA release).

Therefore, we next evaluated mt-dsRNA signatures using normalized A-to-I editing events. Although not significant due to large variability, there was a > twofold increase in mt-dsRNA signatures in AD brains relative to both NCI and MCI groups (Figure 3C). Interestingly, the accumulation of mt-dsRNA in AD brains may be specific to mt-dsRNA because, like we have shown previously^42^, transcriptome-wide dsRNA signatures tended to decrease in AD brains (Figure 3D). Therefore, AD-related changes to mitochondrial maintenance and function may arise from abnormal RNA processing, DNA damage or immune signaling, and it is possible that our analyses actually underestimate the amount of mt-dsRNA present in the AD brain.

To assess the potential functional relevance of AD/mitochondria-related transcriptomic changes, we preformed several co-expression and functional analyses. First, we conducted a weighted gene co-expression network analysis (WGCNA) on protein-coding genes and ChrM transcripts across a subset of ROSMAP samples (n=14) for which we had detailed clinical information (e.g., APOE genotype), neuropathological phenotyping (Braak and CERAD scores), and cognitive performance (memory tests and Mini-Mental State Exam [MMSE]; see Table S3). Most ChrM transcripts were grouped into co-expression module 6 which negatively correlated with Braak score, a clinical measure of neurofibrillary tau tangle distribution in the brain (Figure 3E). Some nuclear-encoded genes important for mitochondrial function also grouped into module 6, and a mitochondrial pathway enrichment analysis revealed that this module was enriched for genes related to metabolism of vitamins D and C, intracellular mitochondrial trafficking and transcription (Figure 3F). A gene ontology enrichment analysis of genes with high module-membership scores (a measure of how similar a gene’s expression profile is to the overall expression pattern of the module) suggested that module 6 genes were relevant to immune signaling (involving TNF and NF-κB), cell proliferation (via ERK/MAPK signaling) and regulating neuronal function/communication (Figure S3B). This could indicate that expression of ChrM transcripts and MitoCarta genes are linked to genes related to neuronal function, proliferation and inflammation.

Two other co-expression modules correlated either positively (module 1) or negatively (module 10) with memory and cognitive function (Figure 3E). Module 1 was enriched for genes related to immune activation (e.g., cytokine and interleukin-1 signaling), metabolism (fatty acid breakdown and glucose homeostasis) and protein modifications, specifically prenylation of CAAX-box proteins, a post-translational modification important for membrane localization including to mitochondrial membranes^43,44^ (Figure S3B). Additionally, there were MitoCarta genes grouped into module 1 enriched for nuclear-encoded mitochondrial genes related to mitochondrial protein import (via TIM23) and biotin-related functions, among others (Figure S3C). Genes grouped in module 10 were related to SCF-KIT signaling (hinting at CNS repair and/or angiogenesis), transcription, and adaptive immunity (Figure S3B). Module 10 was also enriched for nuclear-encoded mitochondrial genes relevant for nucleotide methylation of DNA and rRNAs (via S-Adenosyl-L-methionine [SAM] pathway), protein import, choline metabolism and others that were also enriched in upregulated Age-DEGs (e.g., ROS, glutathione, folate and one-carbon metabolism). These findings suggest that mitochondrial function may be altered in various ways alongside immune activation in individuals with reduced cognitive function as a result of MCI or AD.

### Potential mechanisms of Alzheimer’s disease-associated increases in mt-dsRNA

Despite the trend we observed for AD-related increases in mt-dsRNA (Figure 3C), in our network analyses above (Figure 3E), mt-dsRNA signatures did not correlate with any particular co-expression module. Therefore, to further explore the relationship between mt-dsRNA accumulation, cognitive decline and AD pathogenesis, we examined correlations among mitochondrial transcript abundance (ChrM reads) and mt-dsRNA (A-to-I editing events) with clinical, genetic, and neuropathological features within the ROSMAP clinical subset (Table S3). We found that ChrM expression positively correlated with higher AD risk APOE genotypes (ε4 carriers), indicating that mitochondrial transcriptome regulation may be altered in APOE ε4 carriers in general and could lead to greater mt-dsRNA formation (Figure 3G). Further supporting this possibility, mt-dsRNA signatures were slightly elevated in cognitively normal subjects with higher genetic risk for AD (Figure 3H). Because of these findings, we reanalyzed AD-related changes in gene expression to control for APOE genotype (Figure S4A), and we found that many AD-DEGs related to inflammation and dsRNA signaling (Figure S3A) were no longer upregulated (e.g., MDA5 signaling, defense response to virus; Figure S4B), suggesting that AD-related inflammation/IFN responses may be modulated by an interaction between AD pathology and APOE genotype, and/or that sporadic AD involves downregulation of antiviral defenses. Lastly, upregulated APOE-associated AD-DEGs were associated with ubiquitin-dependent processes (Figure S4B), which is interesting in this context because antiviral signaling and RNA metabolism are regulated by ubiquitination^45–47^. Collectively, these observations suggest that genetic AD risk (i.e., APOE genotype) could impact mt-RNA homeostasis leading to mt-dsRNA accumulation, which could be an unrecognized mechanism that drives genetic susceptibility for the disease.

In this same subset of ROSMAP subjects, mt-dsRNA edits correlated with cognitive function (assessed by mini-mental state exam [MMSE]) and Braak score (Figure 3G), suggesting a clinical-pathological role for mt-dsRNA. To understand what molecular pathways may constitute this link, we performed an intersectional analysis of protein-coding genes that significantly correlated with MMSE, Braak score and/or mt-dsRNA signatures. Genes related to all three measures were involved in DNA binding, cell proliferation, and consistent with previous findings, ubiquitin-dependent protein regulation (Figure 3I). Interestingly, several dsRNA sensors including RIG-1/DDX58, MDA5/IFIH1 are regulated by ubiquitination, and although their expression was only modestly altered in AD (Figure S4C) and did not correlate with mt-dsRNA signatures (Figure 3J), some well-characterized ubiquitin ligases (e.g., TRIM14) and deubiquitinases (e.g., USP47) involved in their regulation were differentially expressed in AD (Figure S4C) and/or correlated with mt-dsRNA signatures in AD brains (Figure 3K). Such changes relevant to regulation of dsRNA metabolism and antiviral/dsRNA signaling in AD, occurring along with age-related mt-dsRNA accumulation, could explain the further increases in mt-dsRNA and greater inflammation in diseased brains.

## DISCUSSION

Neuroinflammation is central to both brain aging and AD^48^. In the aging brain, low-grade chronic inflammation^33^ contributes to synaptic dysfunction, neuronal loss, and cognitive decline. In the AD brain, amyloid-β and tau pathologies further contribute to neuroinflammation, but growing evidence also points to age-related, systemic and cell-intrinsic immune dysregulation as a key factor in disease development^49^. Consequently, current research is increasingly focused on identifying such mechanisms of aging itself that predispose the brain to AD. In line with these efforts, our study identifies mt-dsRNA accumulation as a potential driver of inflammation that arises during aging and is exacerbated by AD, and suggests that dysregulation of the mitochondrial transcriptome may link mechanisms of aging to the development of AD. Our analyses of large transcriptomic datasets revealed that while overall mitochondrial transcript abundance remains relatively stable across the lifespan and with AD, the brains of individuals over 55 years exhibit evidence of significant mt-dsRNA accumulation, which is further elevated in AD brains. Network and co-expression analyses also identified several potential age- and AD-related molecular pathways (discussed below) that may underlie the detected mt-dsRNA accumulation.

Our analyses of transcriptomic changes and mt-dsRNA signatures in aged brains could suggest a model (Figure 4A) in which impaired RNA processing and translation drives mt-dsRNA accumulation. This model idea is supported by existing literature and our data showing: 1) genes involved in mitochondrial RNA processing correlate negatively with mt-dsRNA signatures (i.e., in Figure 2D), and 2) expression of mitochondrial ribosome machinery is reduced (Figure S1B) and correlates positively with mt-dsRNA signatures in older age groups (Figure 2D). These findings suggest that unprocessed and/or untranslated mtRNA may accumulate in aged mitochondria, facilitating spontaneous dsRNA formation like that which occurs in mitochondrial disorders (e.g., Leigh’s syndrome)^50^, various autoinflammatory diseases^51–53^, cancer^54^, and age-related senescence^16^. In older brains, release of mt-dsRNA into the cytoplasm may be enabled by slightly higher expression of non-selective mitochondrial channels/pores such as BAX/BAK channels (Figure S1D) that are known to release mitochondrial RNA and DNA^11,12,55^, which is in consistent data on mt-dsRNA release in senescent cells^16,21^. Once cytoplasmic, mt-dsRNA can activate antiviral signaling proteins (Figure S2B), many of which have elevated expression in older brains (Figure S1C), as others have shown in osteoarthritis, Huntington’s disease and autoimmunity^56^. It has been shown that this mt-dsRNA-induced inflammatory signaling causes DNA damage^57^ and cellular senescence^16^, and may contribute to the development of cancer and neurodegenerative disease. Interestingly, mt-dsRNA accumulation and antiviral signaling proteins (e.g., DDX58/RIG-I and MB21D1/cGAS) were positively correlated in middle-aged brains in our analyses, suggesting that mt-dsRNA-driven inflammation may begin relatively early in aging prior to onset of age-related disease (Figure 2E). Many of these antiviral signaling pathways converge onto the mitochondrial antiviral signaling (MAVS) protein^58^, whose expression increases after ∼35 years of age—the same age at which brain volume begins to decline^59^. Together, these findings support a model (Figure 4A) in which age-related decline in mitochondrial RNA homeostasis causes mt-dsRNA accumulation, release, recognition by cytoplasmic dsRNA sensors, and innate immune activation early in brain aging.

**Figure 4.**
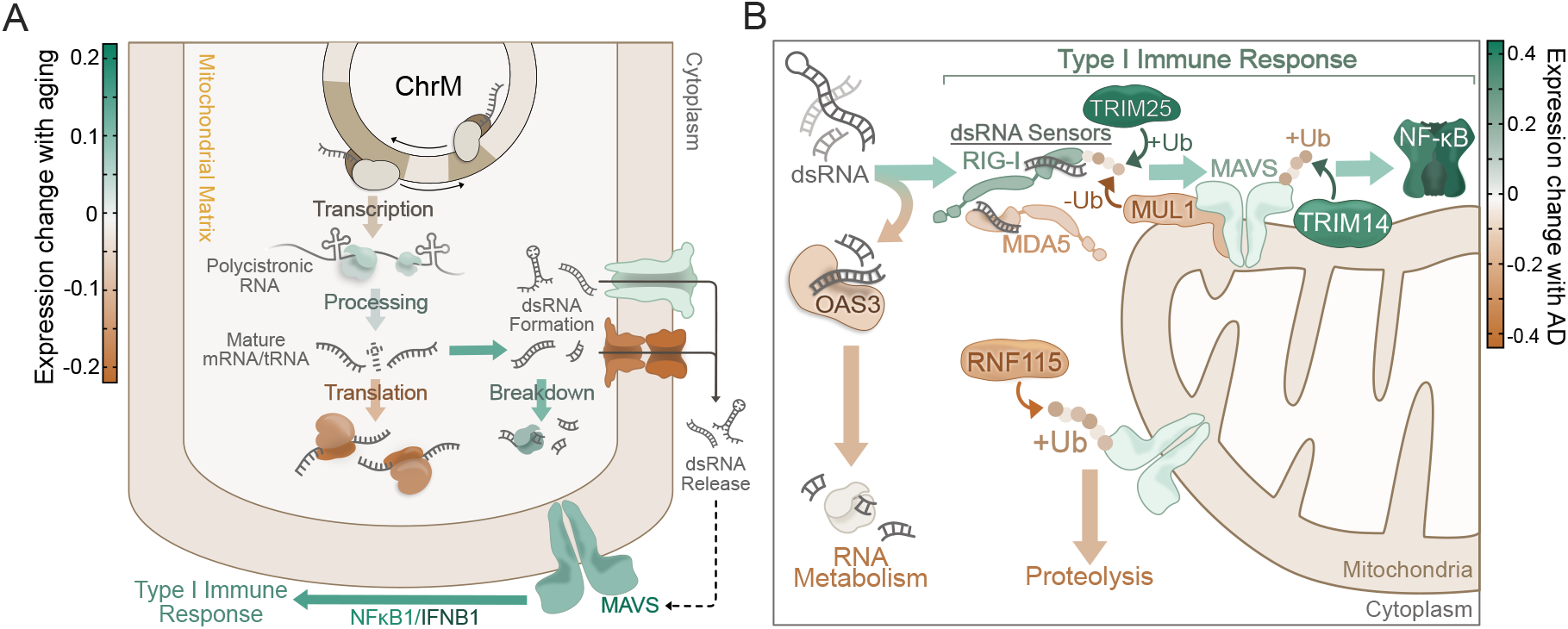
Working model of how the observed changes in mitochondrial and dsRNA signaling proteins may lead to mt-dsRNA accumulation and type I immune activation in aging and AD. A and B) Quantitative illustrations of transcriptomic expression changes (color coded by fold differences) observed in the NABEC (A, for aging) and ROSMAP (B, for AD) datasets. A) In aged brains (>56 y.o. compared to <35 y.o.), expression changes of proteins relevant to mt-dsRNA formation/release and related immune signaling (see Figure S1C) may explain an increase in mt-dsRNA formation and dsRNA-related signaling in older subjects. B) In AD brains, altered expression of dsRNA signaling proteins and their modulators (especially ubiquitin-dependent modulators; see Figure S4C) may decrease dsRNA metabolism and elevate type I immune activation in AD.

As previously mentioned, we also found that age-related mt-dsRNA accumulation may be worsened by AD, and the mechanisms involved may be different from and/or additive to those involved in aging. The AD-related molecular pathways identified in our network and co-expression analyses consistently indicated that ubiquitin-dependent processes are linked to AD pathology, clinical measures of cognitive decline, and mt-dsRNA accumulation. Because RNA metabolism and antiviral signaling are both regulated by ubiquitination, this suggested that ubiquitin-dependent regulation of these processes may be altered with AD and could explain the further increase in mt-dsRNA accumulation in AD brains (see Figure 4B). First, dsRNA metabolism may be hindered in AD due to slight decreases in the dsRNA binding protein OAS3 (Figure S4C), which is responsible for activating the ribonuclease RNase L^60^ that downregulates translation by cleaving rRNA and mRNA to limit viral replication^61^. Also, in our data, expression of OAS3 increases with age (Figure S1C) and was strongly positively related to mt-dsRNA signatures in older, cognitively normal individuals (Figure 3J). This correlation was absent in MCI and slightly reversed in AD brains, even though mitochondrial rRNAs were enriched in AD (Figure 3B) and make up over 20% of the mt-dsRNA detected in our datasets (Figure 2A). This may indicate that depletion of OAS3 in AD allows for accumulation of mitochondrial RNAs, especially rRNAs that readily form dsRNAs.

Second, ubiquitination of antiviral signaling proteins regulates their sensitivity to dsRNA, activity and subcellular localization^45,46^. The results from our functional analyses (Figure 3G) pointed toward changes in ubiquitin-dependent processes in AD brains (Figure 3I and S3A) which may alter the sensitivity of antiviral sensors for dsRNA. For example, RIG-I activation relies on ubiquitination by TRIM25^62^, whose expression was increased by 54% in AD (Figure S4C). Both MDA5 and RIG-I activate MAVS, and MAVS abundance, oligomerization and activation is highly regulated by ubiquitination^45–47^. Their abundance in the mitochondrial outer membrane is attenuated by ubiquitin-dependent degradation of the protein, which is mediated by both mitochondrial-localized and cytoplasmic ubiquitin ligases including MUL1 and RNF115^58^. Expression of both MUL1 and RNF115 was slightly decreased in AD in our analyses (Figure S4C), which would be consistent with lower proteolysis of MAVS, thus increasing abundance and sensitivity of this central dsRNA signaling molecule. At the same time, a different type of ubiquitination (K63-linked) at various sites induces MAVS aggregation/activation and recruits TANK-binding kinase 1 (TBK1), a pivotal kinase in initiating a type I immune response, and TBK1/NF-κB-activating adaptors (TRAFs)^58,63^, which also contribute to tau pathology^64^. Thus, these ubiquitin-dependent regulatory changes may alter the sensitivity of antiviral signaling pathways to endogenous dsRNAs, which could limit appropriate breakdown of cytoplasmic mt-dsRNA and further potentiate chronic inflammation in AD.

In summary, our results suggest that mt-dsRNA represents a novel molecular signal that links mitochondrial dysfunction and innate immune activation in brain aging and AD. This study also identifies a potential mechanism (mt-dsRNA accumulation) by which age-related mitochondrial decline may render the brain susceptible to neurodegeneration and disease progression. Importantly, our conclusions and model are limited, as they are based entirely on retrospective analyses of existing RNA-seq data. Such transcriptomic data, especially from large datasets, can be highly variable and are not directly reflective of protein levels or function, and we were only able to draw inferences about potential underlying mechanisms. Indeed, the purpose of our analyses was not to demonstrate definitive cause and effect, but rather to determine if existing data support the idea of a role for mt-dsRNA in brain aging/AD. To this end, we note that our editing analyses likely detected only a small proportion of mt-dsRNA, the presence of which is dependent on library preparation approaches and may be skewed by related technical artifacts and even single nucleotide variations. As such, our initial exploration of this topic may provide only a small window of insight into the biology and clinical relevance of mt-dsRNA in aging and AD. Future, prospective studies are needed to explore whether mitochondrial RNA homeostasis or mt-dsRNA release are viable targets for treating age- and AD-related neurodegeneration, perhaps using *in vitro* or animal model systems (although directly targeting the multiple dsRNA species released by mitochondria may be highly technically challenging). Regardless, the current data may serve as a basis for investigating novel mitochondria and dsRNA-associated processes that drive brain aging and AD using additional analysis methods (e.g., proteomics) or via primary research methods.

---

**Supplemental Figure 1:** A-D) Additional analyses of the NABEC dataset. A) Combined average expression of genes related to mt-RNA processing, metabolism or translation relative to younger subjects (<35 yrs). Bars represent mean expression of genes (listed in A) relevant to each process ± SEM. **: p_adj_ <0.005 compared to 36-45 y.o. group; Student’s t-test with Bonferroni correction B and C) Expression of nuclear encoded genes involved in B) mt-RNA processing, metabolism and translation, or C) dsRNA sensing and related immune signaling relative to younger subjects (<35 years). *:p<0.05, **:p<0.005 (Student’s t-test; unadjusted for multiple comparisons). D) Relative expression of transcripts encoding mitochondrial pores and channels, and their regulators. *:p_adj_<0.05, **:p_adj_<0.005 (Student’s t-test with Bonferroni correction).

**Supplemental Figure 2:** A) Predicted secondary structure of a single MT-RNR2 transcript (via RNAfold) showing high-probability intrastrand binding. B) Volcano plot of enriched (positive log_2_FC) and depleted (negative log_2_FC) PKR immunoprecipitated transcripts. C-F) Quantification of A-to-I edits in mt-RNA transcripts normalized to ChrM size factor in: C) directly induced into neurons (derived from fibroblasts) originating from younger (30-55 yrs) and older (>56) humans; D) human induced neuron, primary astrocyte and HMC3 microglia co-cultures with and without mitochondrial fission (DRP1) inhibitor (P110); E) vastus lateralis muscle from middle-aged humans (40-64 yrs.) before and after 4 months of daily treatment with 1000 mg Urolithin A; and F) human embryonic lung fibroblasts subject to X-ray induced senescence and/or knock-out (-) of both BAK and BAX genes.

**Supplemental Figure 3:** A-C) Functional analyses related to Figure 3. A and B) Enriched KEGG Reactome, gene ontology biological processes (GO:BP) and molecular function (GO:MF) terms for (A) the top 150 up- or down-regulated AD-DEGs (Figure 3B) and (B) WGCNA gene modules 1, 6 and 10 (Figure 3D). Genes from each module with a module membership greater than 0.9 were included in the gene ontology analysis. C) MitoCarta processes in which module 1 and 10 genes are enriched (using a hypergeometric overlap between gene lists).

**Supplemental Figure 4:** A) Volcano plot of differentially expressed genes in AD compared to NCI subjects (light orange data points, raw p-value < 0.01; dark orange data points indicate genes with FDR < 0.05). B) Enriched KEGG Reactome, GO:BP and GO:MF terms for genes differentially expressed (based on raw p-value < 0.01) in AD individuals when controlling for APOE genotype. C) Quantification of mt-dsRNA signatures grouped by low (e.g., e2/e2 or e2/e3), normal (e3/e3) or high (e3/e4 or e4/e4) genotype within NCI and AD subjects. #: p<0.05, *: p<0.002 (Bonferroni adjusted α).

