## Supplemental File 1 for "Mitochondrial double-stranded RNA accumulation in brain aging and Alzheimer’s disease"

### Evidence of a role for mitochondrial double-stranded RNA accumulation in brain aging and Alzheimer's disease

**Table S1:**

| NABEC Dataset Subject Characteristics | | | |
|---|---|---|---|
| **Age Group** | **n =** | **Sex** | **Age** |
| <35 | 26 | 62% M / 38% F | 30.00 ± 3.84 |
| 36-45 | 36 | 72% M / 28% F | 41.19 ± 2.72 |
| 46-55 | 27 | 66% M / 33% F | 49.74 ± 2.67 |
| >56 | 16 | 81% M / 19% F | 66.81 ± 9.48 |

**Table S2:**

| ROSMAP Dataset Subject Characteristics | | | | | | | | |
|---|---|---|---|---|---|---|---|---|
| **Group** | **n =** | **Sex** | **Age at death** | **Education (total yrs)** | **% with APOE e4** | **MMSE** | **Braak** | **Cerad** |
| NCI | 72 | 75% M 25 % F | 87.60 ± 4.15 | 16.06 ± 3.17 | 16.6% | 27.64 ± 2.53 | 3.19 ± 1.56 | 2.57 ± 1.23 |
| MCI | 38 | 71% M 29% F | 87.95 ± 3.34 | 15.10 ± 2.88 | 7.9% | 24.97 ± 3.64 *** | 3.52 ± 1.31 | 2.07 ± 0.99 |
| AD | 106 | 79% M 21% F | 88.87 ± 2.45 | 15.34 ± 3.09 | 29.2% | 12.06 ± 9.01 *** | 4.34 ± 0.98 *** | 1.66 ± 0.87 *** |

**Table S3:**

| ROSMAP Metadata Subset Subject Characteristics | | | | | | | | | | | | | |
|---|---|---|---|---|---|---|---|---|---|---|---|---|---|
| **Group** | **n =** | **Sex** | **Age at death** | **Educ. (total yrs)** | **MMSE** | **Braak** | **Cerad** | **Global Cognition** | **Episodic Memory** | **Semantic Memory** | **Working Memory** | **Percep. Orientation** | **Speed Percep.** |
| NCI | 3 | 33% M 66 % F | All > 90 | 13.33 ± 1.15 | 28.00 ± 1.00 | 3.00 ± 0 | 3.00 ± 1.00 | 0.04 ± 0.16 | 0.28 ± 0.10 | 0.23 ± 0.52 | 0.04 ± 0.52 | -0.11 ± 0.31 | -0.46 ± 0.71 |
| MCI | 4 | 25% M 75% F | All > 90 | 11.5 ± 1.73 | 25.25 ± 3.20 | 3.50 ± 1.73 | 2.00 ± 0.81 | -0.73 ± 0.64 | -0.39 ± 0.37 | -0.35 ± 0.76 | -0.56 ± 0.24 | -0.67 ± 0.49 | -1.14 ± 1.14 |
| AD | 7 | 43% M 57% F | 89.83 ± 0.30 | 14.42 ± 1.98 | 10.43 ± 8.71* | 1.11 ± 0.09 | 2.00 ± 0.81 | -1.58 ± 0.61* | -1.71 ± 0.91* | -1.49 ± 0.88* | -1.02 ± 0.52* | -0.73 ± 0.94 | -1.91 ± 0.38* |
